# Cover crop mixture diversity, biomass productivity, weed suppression, and stability

**DOI:** 10.1101/440180

**Authors:** A. M. Florence, L. G. Higley, R. A. Drijber, C. A. Francis, J. L. Lindquist

**Affiliations:** Department of Agronomy and Horticulture, University of Nebraska-Lincoln, Lincoln, Nebraska, United States of America; School of Natural Resources, University of Nebraska-Lincoln, Lincoln, Nebraska, United States of America

## Abstract

The diversity-productivity, diversity-invasibility, and diversity-stability hypotheses propose that increasing species diversity should lead, respectively, to increased average biomass productivity, increased invasion resistance, and increased stability. We tested these three hypotheses in the context of cover crop mixtures, evaluating the effects of increasing cover crop mixture diversity on aboveground biomass, weed suppression, and biomass stability. Twenty to forty cover crop treatments were replicated three or four times at eleven sites using eighteen species representing three cover crop species each from six pre-defined functional groups: cool-season grasses, cool-season legumes, cool-season brassicas, warm-season grasses, warm-season legumes, and warm-season broadleaves. Each species was planted in monoculture, and the most diverse treatment contained all eighteen species. Remaining treatments included treatments representing intermediate levels of cover crop species and functional richness and a no cover crop control. Cover crop planting dates ranged from late July to late September with both cover crop and weed aboveground biomass being sampled prior to winterkill. Stability was assessed by evaluating the variability in cover crop biomass for each treatment across plots within each site. While increasing cover crop mixture diversity was associated with increased average aboveground biomass, this was the result of the average biomass of the monocultures being drawn down by low yielding species rather than due to niche complementarity or increased resource use efficiency. At no site did the highest yielding mixture out-yield the highest yielding monoculture. Furthermore, while increases in cover crop mixture diversity were correlated with increases in weed suppression and increases in biomass stability, we argue that this was largely the result of diversity co-varying with aboveground biomass, and that differences in aboveground biomass rather than differences in diversity drove the differences observed in weed suppression and stability. The results of this study contradict popular interpretations of the diversity-productivity, diversity-invasibility, and diversity-stability hypotheses.

## Introduction

Increasing species diversity is thought to lead to increased average productivity, increased invasion resistance, and increased stability [1, 2]. Respectively named the diversity-productivity, diversity-invasibility, and diversity-stability hypotheses, these three hypotheses, while contested in the field of ecology [3–5], have often been treated in the field of agriculture as proven principle with regard to mixed cropping. Increasing crop mixture diversity is often assumed to be associated with increased productivity, increased weed suppression, and increased stability despite a lack of compelling empirical evidence in favor of these assertions [6–9]. The goal of this study is to test these three hypotheses in the context of cover crop mixtures.

Cover crops are used to provide a variety of functions, many of which are positively related to cover crop productivity. These functions include weed suppression, soil nutrient retention, soil erosion control, and organic matter addition. While the use of cover crops in agriculture has a long history, the use of highly diverse cover crop mixtures is a relatively recent phenomenon. It has been suggested that by increasing cover crop mixture diversity, the various functions of cover crops will be enhanced and stabilized. Specifically, it has been proposed in both the popular press and the scientific literature that increasing cover crop mixture diversity should be associated with increased productivity, increased weed suppression, and increased biomass stability—claims that parallel the assertions made by the diversity-productivity, diversity-invasibility, and diversity-stability hypotheses (e.g. [10–14]).

While the diversity-productivity, diversity-invasibility, and diversity-stability hypotheses may appear to address three distinct topics, niche differentiation between species is used as the logical basis for all three. Niche differentiation implies that different species have different resource needs and acquisition abilities. Consequently, a single species is expected to leave resources unexploited that another species might be able to exploit—e.g., through its differential root or canopy architecture. Thus, the diversity-productivity hypothesis expects that a more diverse system should be more productive than a less diverse system due to increased resource use efficiency or niche complementarity [15]. Also, since a more diverse community is expected to more fully use the finite resources in an environment than a less diverse community, the diversity-invasibility hypothesis predicts that a more diverse community will also be less susceptible to invasion by other species than a less diverse community, as more of the available resources have been pre-empted [5]. Furthermore, since different species have different resource requirements and physiological efficiencies, it follows that different species will thrive and fail under different conditions. Consequently, the diversity-stability hypothesis predicts that the presence of many species insures that at least some species will thrive under variable environmental conditions, thereby stabilizing the performance of the species mixture [4].

Despite the apparent logical soundness of these hypotheses, there is limited evidence supporting them, particularly in the context of agriculture. Using cover crop mixtures as our model system, we ask with this study: Does increasing cover crop mixture diversity (1) increase cover crop biomass productivity, (2) increase weed suppression, and/or (3) increase biomass stability?

## Materials and methods

### Research sites

This study was conducted at eleven sites on practicing farms across southeastern Nebraska. Cover crops were sown at a variety of times in a variety of crop rotations (Table 1). With the exception of sites 1 and 4, where the farm was irrigated, all other sites were rain-fed.

**Table 1.** Study locations, planting dates, planting conditions, and sampling dates. In cases where cover crops were seeded into a maturing crop, the growth stage of that crop is also provided in parentheses.

| Site | Location | Cover crop planting date | Planting conditions | Sampling date* |
| --- | --- | --- | --- | --- |
| 1 | 40°24'60"N 99° 2'60"W | 7/19/2013 | Wheat stubble | - |
| 2 | 40°58'25"N 97°59'15"W | 8/10/2013 | Barley stubble | - |
| 3 | 41°40'15"N 96°33'45"W | 8/31/2013 | Wheat stubble (disked) | 10/31/2013 |
| 4 | 41°10'20"N 96°27'30"W | 9/10/2013 | Soybeans (R5) | 11/9/2013 |
| 5 | 41°40'10"N 96°33'50"W | 9/12/2013 | Soybeans (R7) | 11/7/2013 |
| 6 | 41°40'20"N 96°34'5"W | 9/12/2013 | Corn (R6) | - |
| 7 | 40°58'10"N 97°59'50"W | 9/14/2013 | Soybeans (R6) | 11/14/2013 |
| 8 | 41°19'45"N 96°16'55"W | 9/19/2013 | Corn stubble (disked) | 11/8/2013 |
| 9 | 40°19'5"N 98°35'45"W | 9/20/2013 | Corn (R6) | - |
| 10 | 41°40'20"N 96°33'40"W | 7/20/2014 | Wheat stubble (disked) | 9/27/2014 |
| 11 | 40°51'5"N 96°28'10"W | 7/23/2014 | Wheat stubble | 10/14-15/2014 |

\*Not all sites were sampled for plant biomass.

### Treatments

The study was started in 2013 with twenty treatments representing monocultures and mixtures of nine species—barley, oat, wheat, Austrian winter pea, red clover, yellow sweetclover, radish, rapeseed, and turnip (Table 2). The nine species were selected to represent three functional groups—cool-season grasses, legumes, and brassicas. The grasses used were all spring varieties, which winterkilled along with the legumes and brassicas.

**Table 2.** Summary of cover crop treatments for 2013.

| No. | Functional group(s) | Treatment | No. of species | No. of groups |
| --- | --- | --- | --- | --- |
| 1 | - | No cover | 0 | 0 |
| 2 | Cool-season grasses (CG) | Barley (BAR) | 1 | 1 |
| 3 |  | Oats (OAT) | 1 | 1 |
| 4 |  | Wheat (WHT) | 1 | 1 |
| 5 | Cool-season legumes (CL) | Austrian winter pea (PEA) | 1 | 1 |
| 6 |  | Red clover (RED) | 1 | 1 |
| 7 |  | Yellow sweetclover (YEL) | 1 | 1 |
| 8 | Cool-season brassicas (CB) | Radish (RAD) | 1 | 1 |
| 9 |  | Rapeseed (RAPE) | 1 | 1 |
| 10 |  | Turnip (TURN) | 1 | 1 |
| 11 | CG | BAR + OAT + WHT | 3 | 1 |
| 12 | CL | PEA + RED + YEL | 3 | 1 |
| 13 | CB | RAD + RAPE + TURN | 3 | 1 |
| 14 | CG + CL | BAR + OAT + WHT + PEA + RED + YEL | 6 | 2 |
| 15 | CG + CB | BAR + OAT + WHT + RAD + RAPE + TURN | 6 | 2 |
| 16 | CL + CB | PEA + RED + YEL + RAD + RAPE + TURN | 6 | 2 |
| 17 | CG + CL + CB | All 9 cool-season species | 9 | 3 |
| 18 | CG + CL + CB | BAR + PEA + RAD | 3 | 3 |
| 19 |  | OAT + RED + RAPE | 3 | 3 |
| 20 |  | WHT + YEL + TURN | 3 | 3 |

Treatment 1 was a no cover control. Treatments 2-10 were all the species included in the study grown in monoculture. Treatments 11-13 were mixtures of all three cool-season grasses, legumes, and brassicas, respectively. These treatments served to evaluate the effect of increasing species diversity without increasing functional diversity.

Treatment 14 combined the grasses with the legumes, treatment 15 combined the legumes with the brassicas, and treatment 16 combined the grasses with the brassicas. These three treatments served as a level of functional diversity intermediate between treatments 11, 12, and 13 and treatment 17, which combined all nine species used.

Treatments 18-20 were combinations of one grass, one legume, and one brassica. These last three treatments were designed so that each of the nine species was present in one of the three treatments. In designing all of the treatments used, a point was made to make sure that each species was equally represented at each level of species and functional richness to address the issue of sampling bias—that is, the issue that as diversity increases, the likelihood of a certain species being included also increases [16–18]. Beyond that criteria being met, the specific combination of each grass, legume, and brassica was arbitrary.

In 2014, the study was expanded to include an additional 20 treatments (Table 3). Of these additional treatments, treatments 21-39 represented warm-season analogues of treatments 2-20. That is, warm-season grasses, legumes, and broadleaves were used instead of the cool-season grasses, legumes, and brassicas. The species used were proso millet, sorghum sudangrass, teff, chickpea, cowpea, sunn hemp, buckwheat, safflower, and sunflower. Treatment 40 was a combination of the original nine cool-season species and these nine warm-season species. For a discussion of the traits associated with the cover crop species used in this study, refer to Clark [19].

**Table 3.**
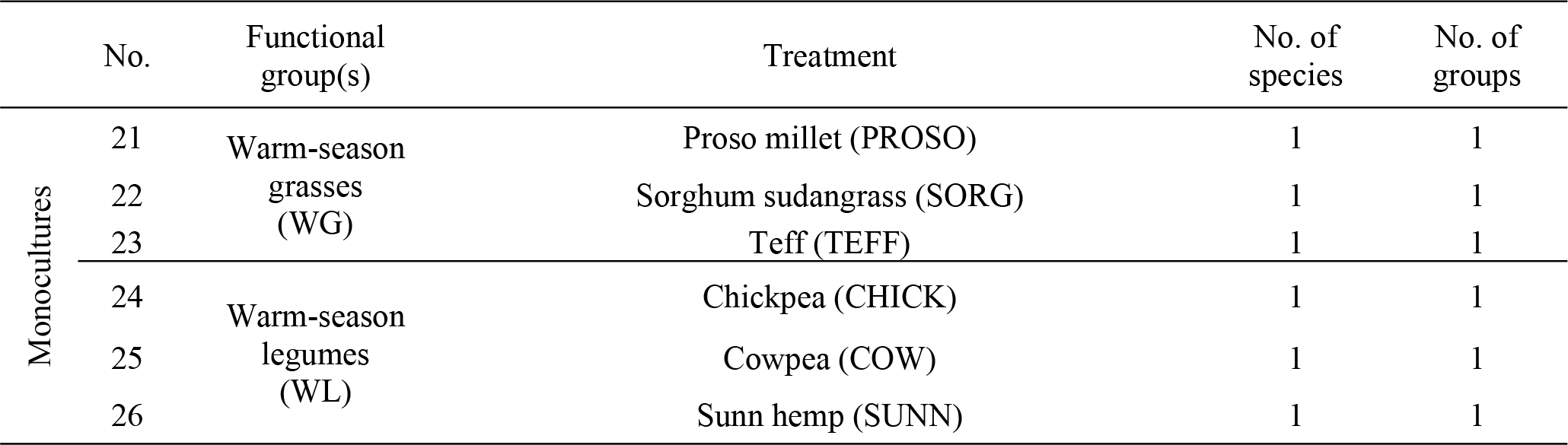
Summary of cover crop treatments added in 2014.

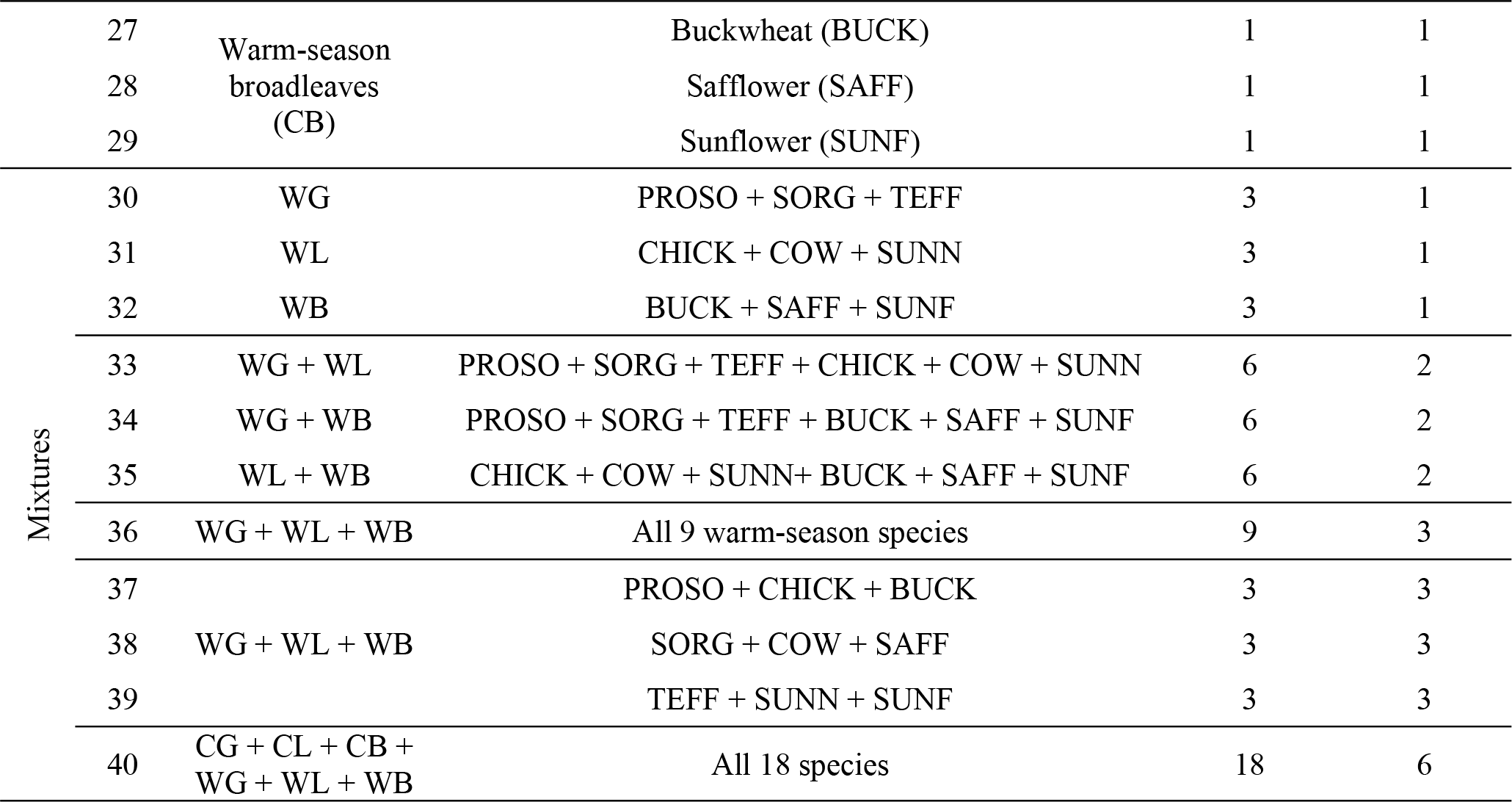

### Seeding rates

Seeding rates for the different cover crops in monoculture were based on recommended broadcast rates [19] (Table 4). Cover crop mixture seeding rates were proportional to the rates used in monoculture. For example, in a three species mix, each species was planted at one-third the full rate. The seeding rates for the brassica species were reduced in the second year of this study as the original seeding rate was greater than necessary to achieve maximum biomass.

**Table 4.**
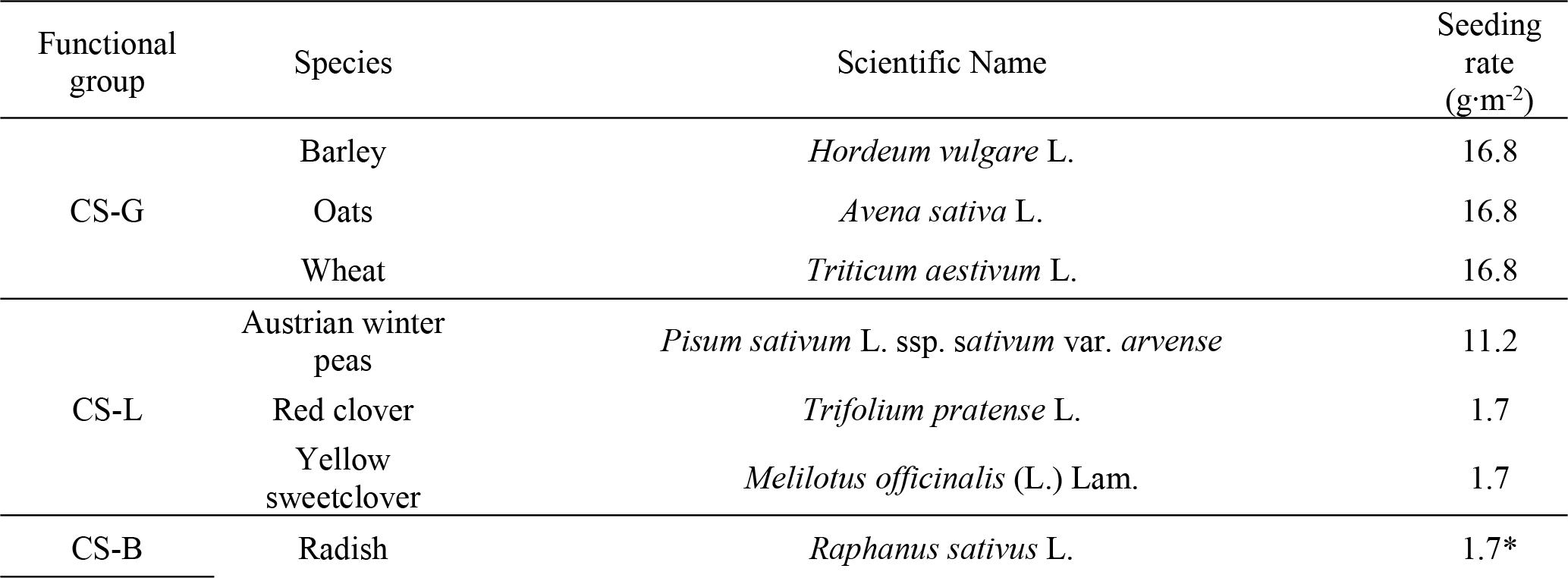
Seeding rates used for each cover crop species in monoculture.

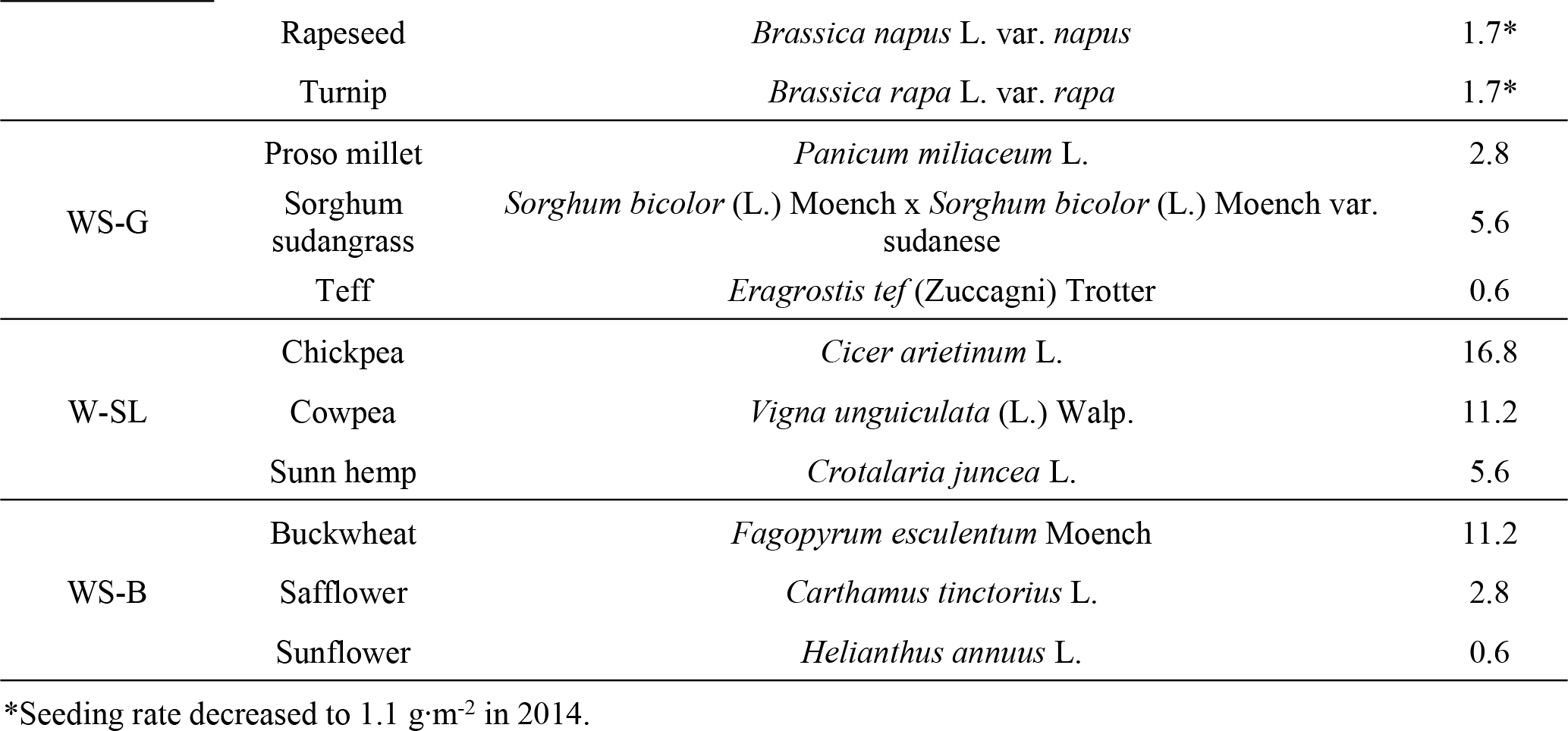

### Treatment establishment

Treatments were arranged in a randomized complete block design with four replications at each site with the exception of site 11, which had only three replications owing to space constraints. Plots were 5 m x 10 m—though these dimensions varied slightly to accommodate corn and soybean row spacing at sites 4, 5, 6, 7, and 9. Seeds for each treatment were hand broadcast into a variety of field conditions—after small grains harvest, after corn harvest, and into maturing corn and soybeans. In some instances, harvested small grain fields were disked prior to cover crop seeding and establishment. In other instances, the cover crop seeds were broadcast into standing stubble (Table 1). Field management decisions were left up to each cooperating farmer.

### Data collection

Cover crop aboveground biomass was harvested prior to winterkill. Where sufficient growth was present (sites 3, 10, and 11), weed aboveground biomass was also sampled at this time. Vegetation was sampled using two randomly placed 0.18 m^2^ quadrats in each plot for site 3 and one randomly placed 0.18 m^2^ quadrat in each plot for the rest of the sites harvested. For perspective, many planted diversity studies use a sample of 0.20 m^2^ per plot [20]. Cover crop biomass was separated to species. Weed biomass was also separated to species with the exception of *Amaranthus spp*. and *Setaria spp*., which were separated to genus. Plant samples were dried at 55°C for 7 days and dry mass determined.

### Data analysis

#### Diversity-productivity hypothesis

The diversity-productivity hypothesis was tested by calculating estimates of the effect size of increasing species and functional richness on biomass productivity.

To separate the effects of species richness from the effects of functional richness, we asked the question: “Does increasing species richness without increasing functional richness increase aboveground biomass?” We approached this question in two ways: (1) by tripling species richness within each functional group, and (2) by tripling the species richness of already functionally diverse mixtures. In the first case, for example, the difference between the biomass of the 3 species all grass mixture (treatment 11) and the average biomass of the constituent grass monocultures (treatments 2, 3, and 4—barley, oats, and wheat, respectively) was divided by the latter. This was also done for the 3 species all legume and all brassica mixtures (treatment 12 and 13, respectively).

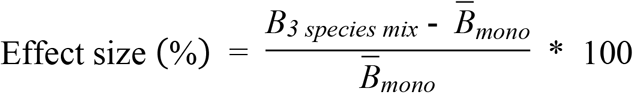

In the second case, we compared the average aboveground biomass of treatments containing one cool-season grass, legume, and brassica (*B̅*_*18,19,20*_) with treatment 17, which contained three cool-season grasses, three cool-season legumes, and three brassicas (*B*_*17*_).

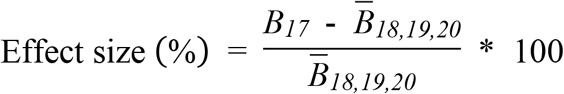

To determine the effect of increasing functional richness alone, we held species richness constant at three species and increased functional richness from one functional group to three. That is, we compared the aboveground biomass of treatments 11, 12, and 13 to treatments 18, 19, and 20.

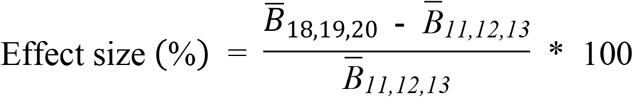

The effect of increasing species richness and functional richness simultaneously was tested by taking the aboveground biomass of the nine-species mixture (i.e., treatment 17) and subtracting the average aboveground biomass of those nine species in monoculture (i.e., treatments 2-10), and then dividing by the average production of the monocultures.

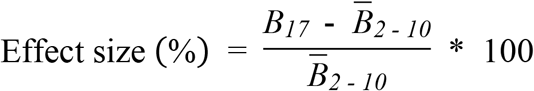

Calculating this value for each block at each site results in multiple estimates of effect size. To these approximately normal populations of estimates, we applied simple one-sample t-tests to determine the effects of (1) increasing species richness alone, (2) increasing functional richness alone, and (3) increasing species and functional richness together. Due to irregularities in the warm-season species data, which will be discussed in the results, as well as the low number of replicates of these treatments, these treatments were excluded from this analysis, though treatment summary data are provided.

#### Diversity-invasibility hypothesis

The diversity-invasibility hypothesis was tested by evaluating whether increasing cover crop diversity increased weed suppression of a cover crop on a per unit biomass basis (Fig 1a). To test this hypothesis, we first calculated percent weed biomass reduction (*BR*_*weed*_) as:

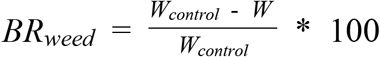

Where *w*_*control*_ is the average weed biomass in the control (no cover crop) plots for each site and *w* is the weed biomass in each cover crop plot. Then, *BR*_*weed*_ was related to cover crop biomass (*x*) by an exponential equation:

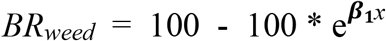

Where *β*_*1*_ is a fitted parameter indicating the responsiveness of weed biomass to cover crop biomass—the larger the *β*_*1*_ parameter, the more responsive weed biomass is to cover crop biomass. To assess whether species richness affects invasibility after controlling for the effect of cover crop biomass, a modified version of the equation was also fit:

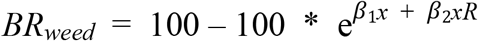

Where *R* was either cover crop species richness or functional richness—as measured by the number of cover crop species or functional groups identified in the sampling quadrat—and *β*_*2*_ was an additional fitted parameter that allows for cover crop diversity to affect the relationship between percent weed biomass reduction and cover crop biomass. The significance of the parameter estimate *β*_*2*_, based on an F-test, was used to draw conclusions about the impact of species richness and functional richness on invasibility.

**Fig 1.**
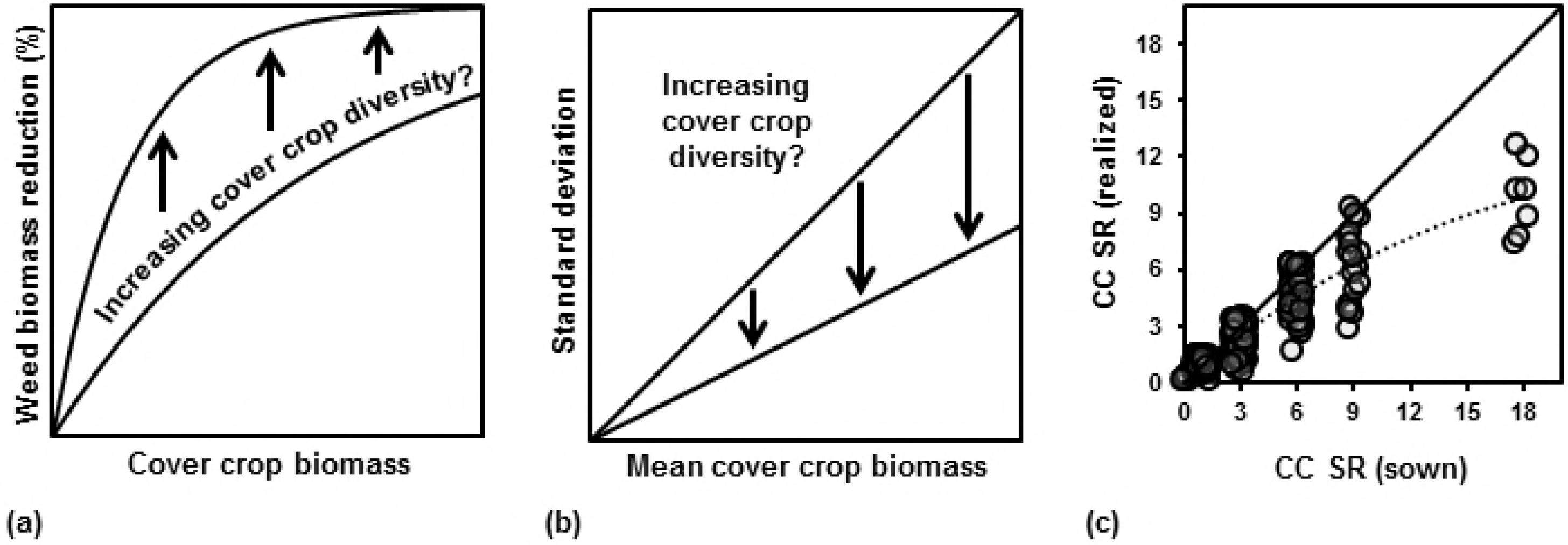
Hypothesized effect of species diversity on invasibility and stability. (a) effect of increasing cover crop diversity (species or functional richness) on the relationship between weed biomass reduction and cover crop biomass. (b) effect of increasing cover crop diversity on the relationship between standard deviation of cover crop biomass and mean cover crop biomass. (c) Realized cover crop species richness versus planted cover crop species richness. Points jittered along both axes for ease of viewing. Solid line indicates an idealized 1:1 relationship. Dashed line indicates LOESS curve fitted to data (α=1, λ= 2).

#### Diversity-stability hypothesis

The term “stability” is used in the ecological and agricultural literature to refer to different ideas and is consequently measured in different ways [21]. The standard metric to evaluate stability is the coefficient of variation (C_v_) of stand biomass, which is estimated as the sample standard deviation of the mean biomass (s) divided by the sample mean biomass (x̅). A low C_v_ is considered an indicator of high stability and a high C_v_ an indicator of low stability. Generally, the C_v_ is then regressed on a diversity metric like species richness [22], with a negative slope indicating increased stability with increasing diversity. However, the results of this analysis can be misleading because the effects of diversity on stability are confounded with the effects of biomass productivity on stability. To avoid these confounding effects, we selected a metric that would indicate how consistent stand biomass was from plot to plot, and regressed the standard deviation of cover crop biomass against mean cover crop biomass [23] for each treatment at each site and tested whether increasing cover crop diversity—as measured by cover crop species and functional richness—decreased the slope of this relationship (**Error! Reference source not found.**). In essence, this assesses whether plot to plot variability within a field decreased with increasing cover crop diversity.

### Sown versus realized species richness

In diversity studies looking at plant mixtures, authors often have to make a decision of whether to look at sown diversity—how many species or functional groups were planted—or realized diversity—how many species or functional groups actually germinated and survived, and thus were observed. Realized diversity typically correlates well to sown diversity but the deviation between realized and sown species richness tends to increase with increasing sown species richness (**Error! Reference source not found.**).

When evaluating the effect of cover crop mixture diversity on weed suppression, we judged that realized diversity was the more appropriate metric to use—as any species or functional group that was sown but absent in our sampling was unlikely to have an effect on the weed biomass in our sampling. However, using sown diversity values instead of realized diversity values resulted in the same interpretive conclusions.

When evaluating the effect of diversity on stability, we judged that sown diversity was the more appropriate metric to use—as the diversity-stability hypothesis is predicated on the idea that a more species-rich mixture is better insured against the failure of any one species. However, using realized diversity instead of sown species richness also resulted in the same interpretive conclusions.

### Statistical software

All statistical analyses were conducted using R 3.1.0 [24]. Non-linear regression models were fit with the nls2 package by Grothendieck [25].

## Results

### Cover crop productivity by site

Cover crops were not harvested at 4 of the 11 sites planted. At site 1, cover crop establishment was patchy throughout the site due to wheat stubble being swathed after cover crop planting. At site 2, there was negligible cover crop growth due to extreme weed pressure. At sites 6 and 9 there was negligible cover crop growth (< 25 g m^−2^)— likely due to a combination of limited water and light under the standing corn crop and heat stress. Of those sites that were harvested, the earlier planting dates had the greatest aboveground biomass, with negligible biomass for those sites planted after the beginning of September (Fig 2).

**Fig 2.**
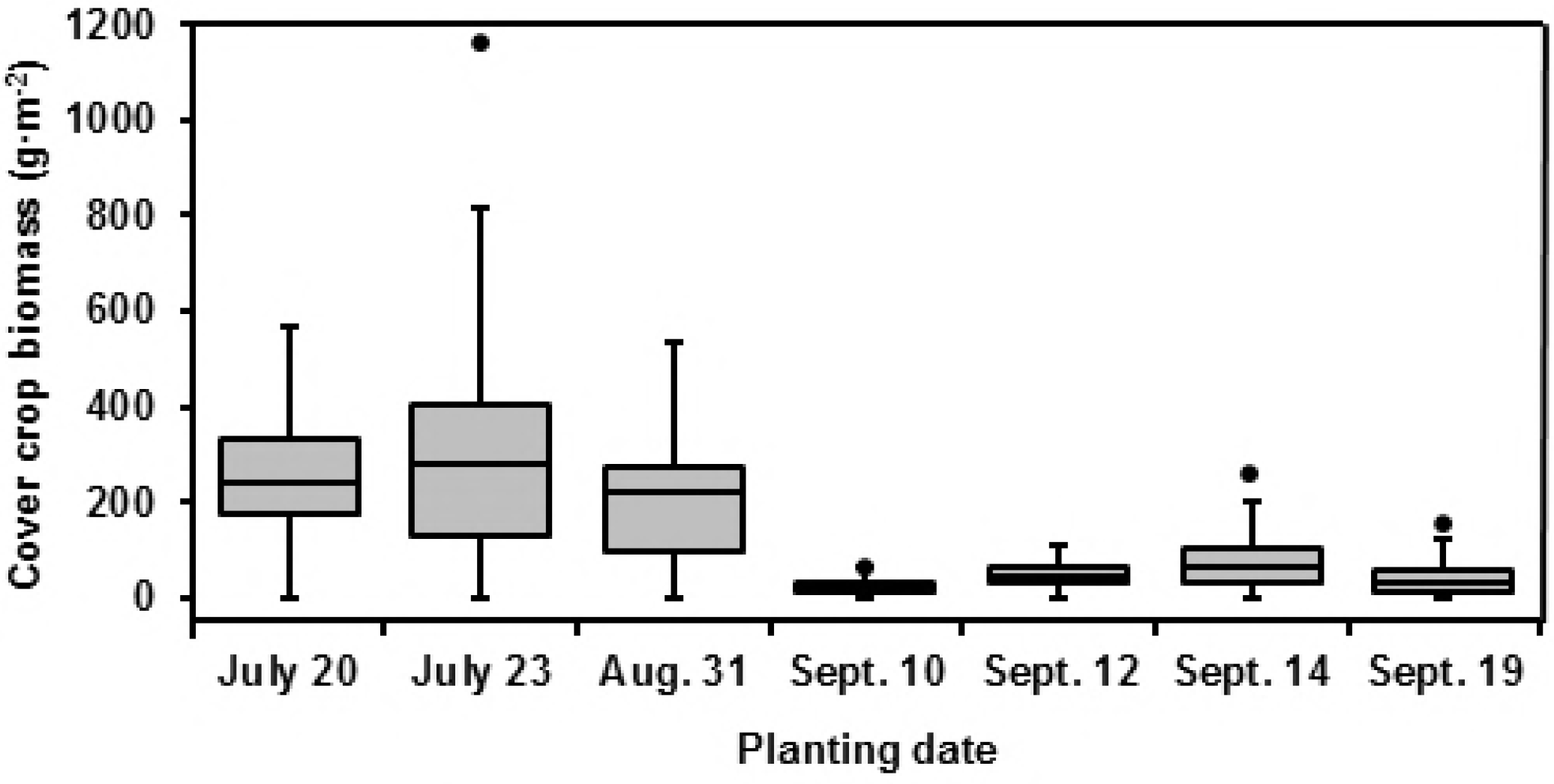
Cover crop productivity and time of establishment. Boxplots of cover crop aboveground biomass for treatments #2-20 by planting date. Planting dates are not temporally equidistant.

### Cover crop productivity by treatment

Cover crop productivity by treatment varied widely across sites, but a few patterns were consistent across all sites. Cool-season grasses and brassicas generally out-produced the legumes (Figs 3 and 4) with the best performing species varying among sites. Warm-season grasses tended to out-produce the legumes (Fig 5). The relatively poor performance of the legumes may have been related to the relatively nutrient rich condition of the sites used. Buckwheat was consistently one of the most productive warm-season broadleaf species, safflower was one of the least productive, and sunflower productivity was inconsistent across sites. This is likely due to deer having grazed on the sunflower plants prior to sampling at site 11 but not site 3. Sampling at sites 3 and 11 happened after some of the warm-season species began to shed their foliage, leading to lower measured aboveground biomass than was actually produced. These irregularities in the warm-season species should be kept in mind when considering the results.

**Fig 3.**
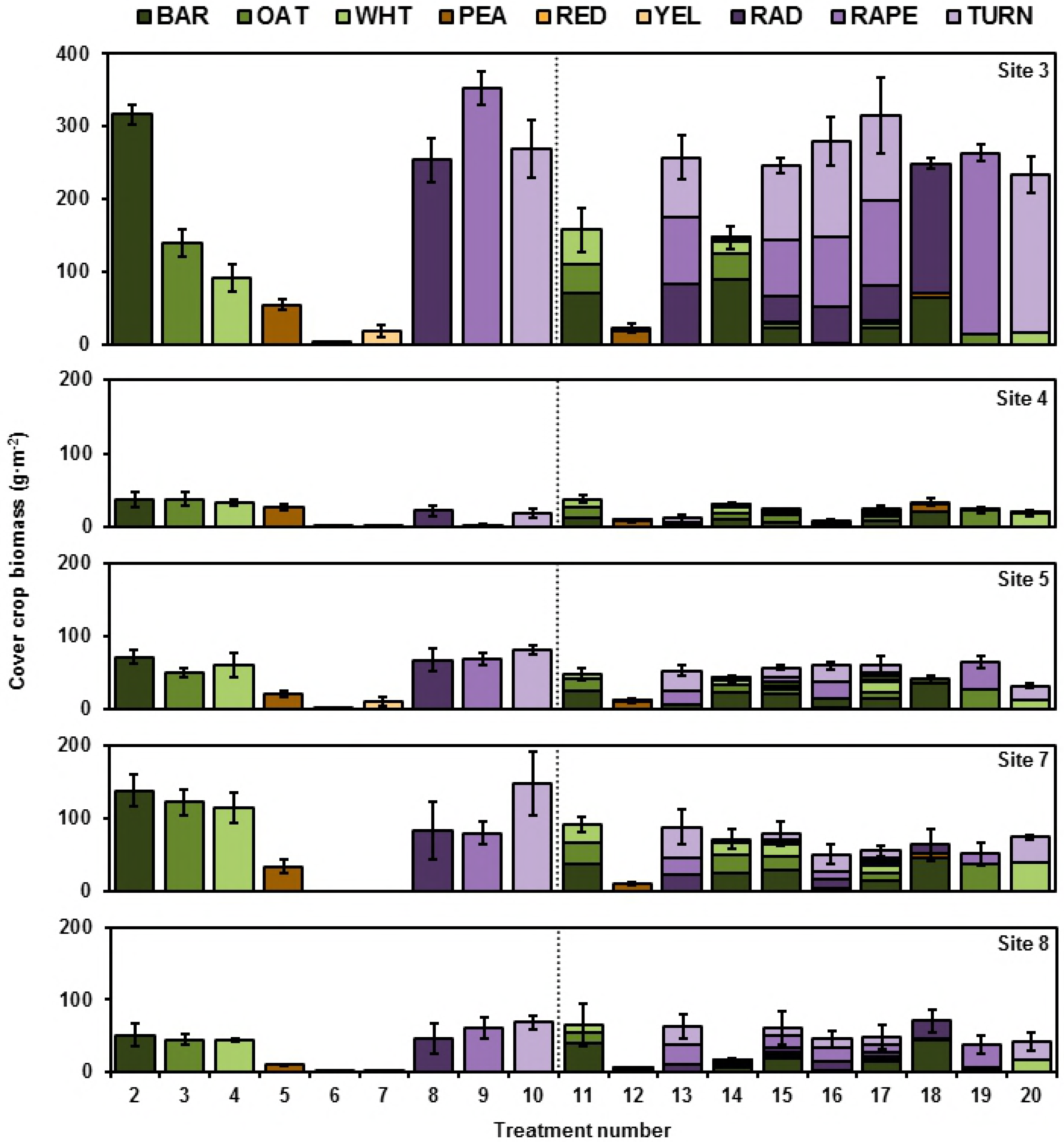
Cover crop productivity 2013. Species specific cover crop biomass (±SEM) for treatments 2-20 by site for 2013. The vertical dotted line separates monoculture (left) from mixtures (right). BAR = barley. OAT = oat. WHT = wheat. PEA = Austrian winter pea. RED = Red clover. YEL = Yellow sweetclover. RAD = Radish. PARE = Rapeseed. TURN = turnip.

**Fig 4.**
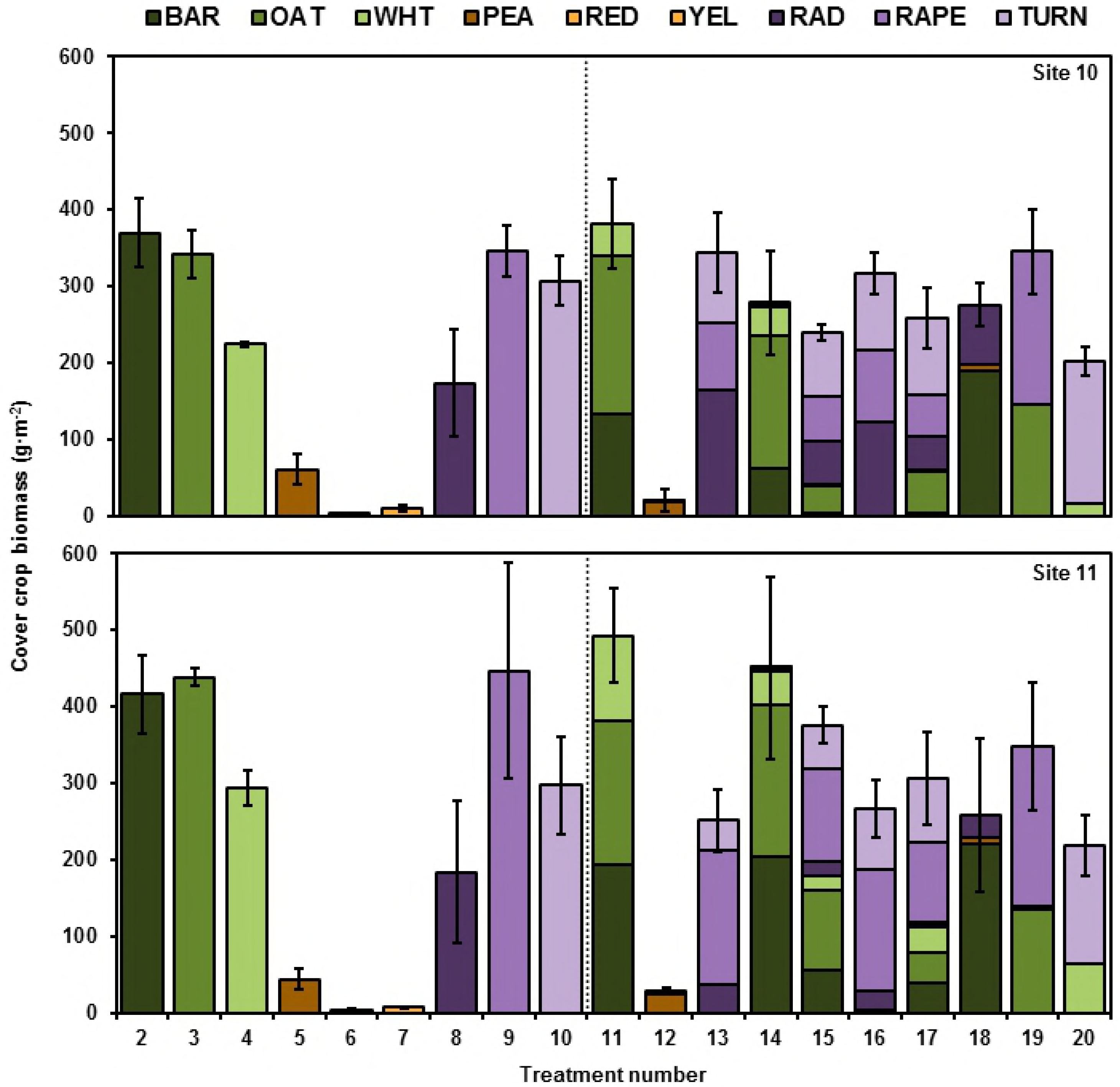
Cool season cover crop productivity 2014. Species specific cover crop biomass (±SEM) for treatments 2-20 by site for 2014. The vertical dotted line separates monoculture (left) from mixtures (right). One extreme outlier (1156 g·m^2^) for rapeseed was omitted from the bar chart for Site 11. BAR = barley. OAT = oat. WHT = wheat. PEA = Austrian winter pea. RED = Red clover. YEL = Yellow sweetclover. RAD = Radish. PARE = Rapeseed. TURN = turnip.

**Fig 5.**
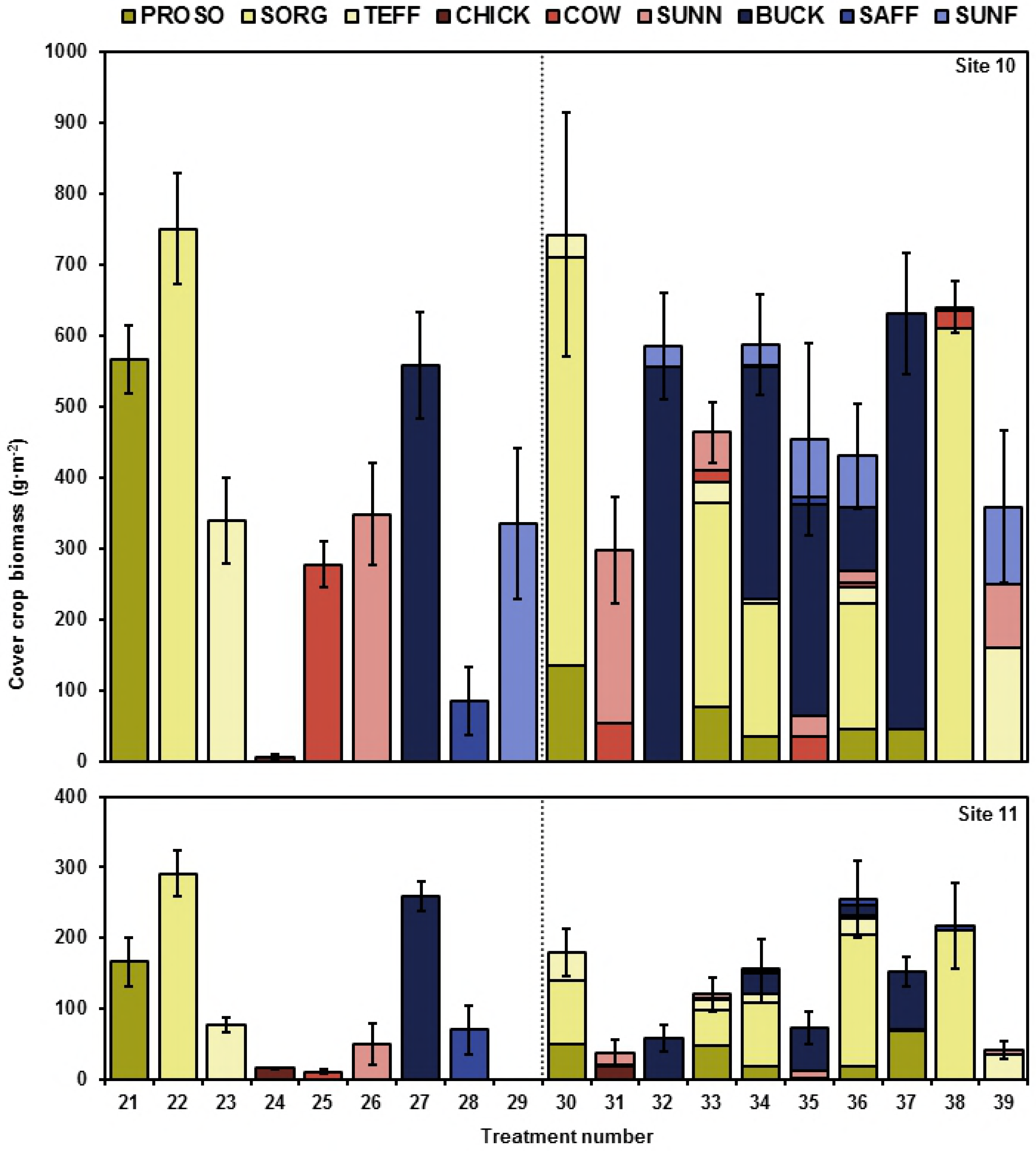
Warm season cover crop productivity 2014. Species-specific cover crop biomass (±SEM) for treatments 21-39 by site for 2014. The vertical dotted line separates monocultures (left) from mixtures (right). PROSO = proso millet. SORG = Sorghum sudangrass. TEFF = teff. CHICK = chickpea. COW = cowpea. SUNN = sunn hemp. BUCK = buckwheat. SAFF = safflower. SUNF = sunflower.

Cool-season mixtures tended to be dominated by brassicas and warm-season mixtures tended to be dominated by sorghum sudangrass and buckwheat, when present. A species’ biomass in monoculture was fairly predictive of its biomass in mixture, such that high-yielding species in monoculture were also high yielding in mixture and vice versa.

### Effect of diversity on biomass productivity

Increasing species richness, while holding functional richness constant, did not increase average aboveground biomass (mean effect size = 2.3%, 95% C.I. = [−7.2, 11.9%], N = 107, *p*-value = 0.65). However, increasing functional richness, while holding species richness constant, increased average aboveground biomass by 29%, and increasing both functional and species richness simultaneously increased average aboveground biomass by 28% (Fig 6). Note, however, that at no site did any mixture out-yield the most productive monoculture (Figs 3-5).

**Fig 6.**
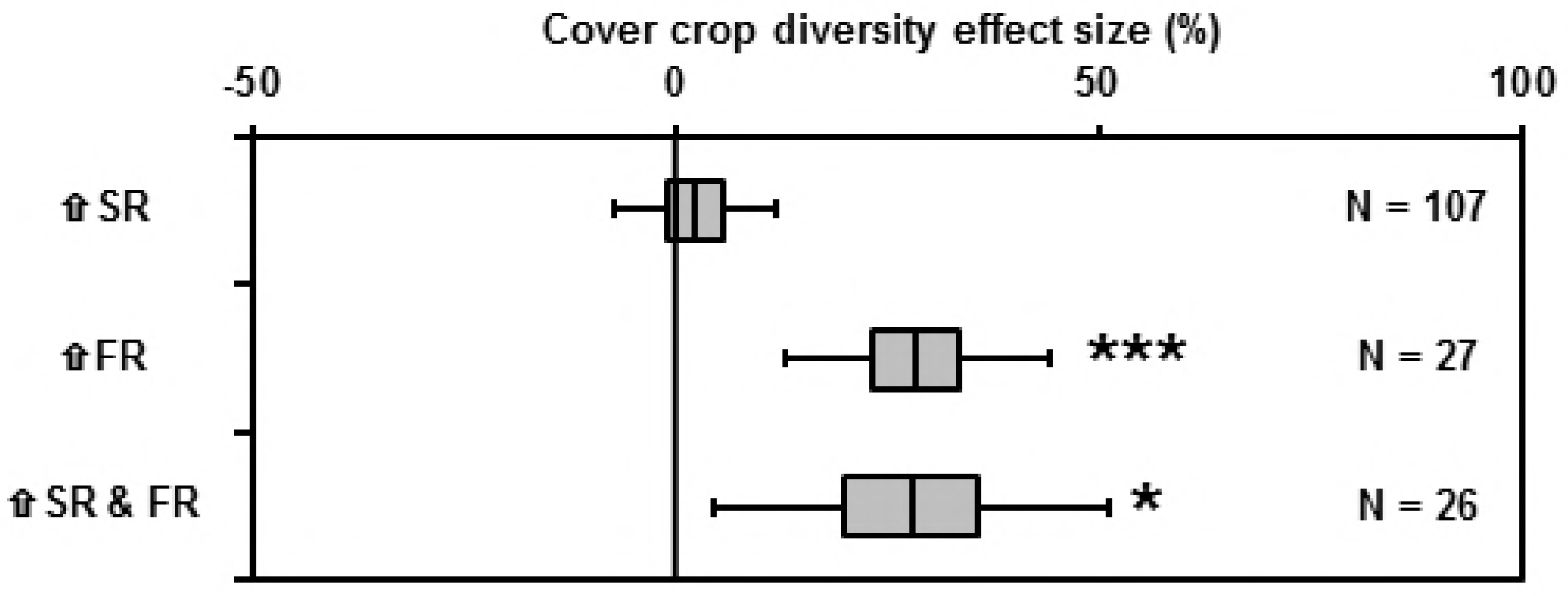
Effect of diversity on biomass productivity. Mean effect size of increasing cover crop diversity on cover crop productivity—specifically the effects of increasing species richness (⇧SR), increasing functional richness (⇧FR), and increasing both species and functional richness simultaneously (⇧SR & FR). Boxes and bars represent 50% and 95% confidence intervals, respectively. N = number of observations for each estimate. One observation is missing from the ⇧SR & FR category. Asterisks indicate *p*-value for the following test—H_0_: μ = 0; H_a_: μ ≠ 0. *P*-value > 0.05 (no asterisk); < 0.05(*); < 0.01(**); < 0.001(***).

### Effect of diversity on weed suppression

Increased cover crop biomass was associated with increased weed suppression at all three sites (Fig 7). However, neither adding cover crop species richness nor functional richness values improved the predictive results of the models tested with the exception of adding functional richness to the site 10 base model, which resulted in a marginal improvement in predictive results (Table 5). Overall, the impression given is that increasing cover crop mixture diversity did not increase weed suppression.

**Fig 7.**
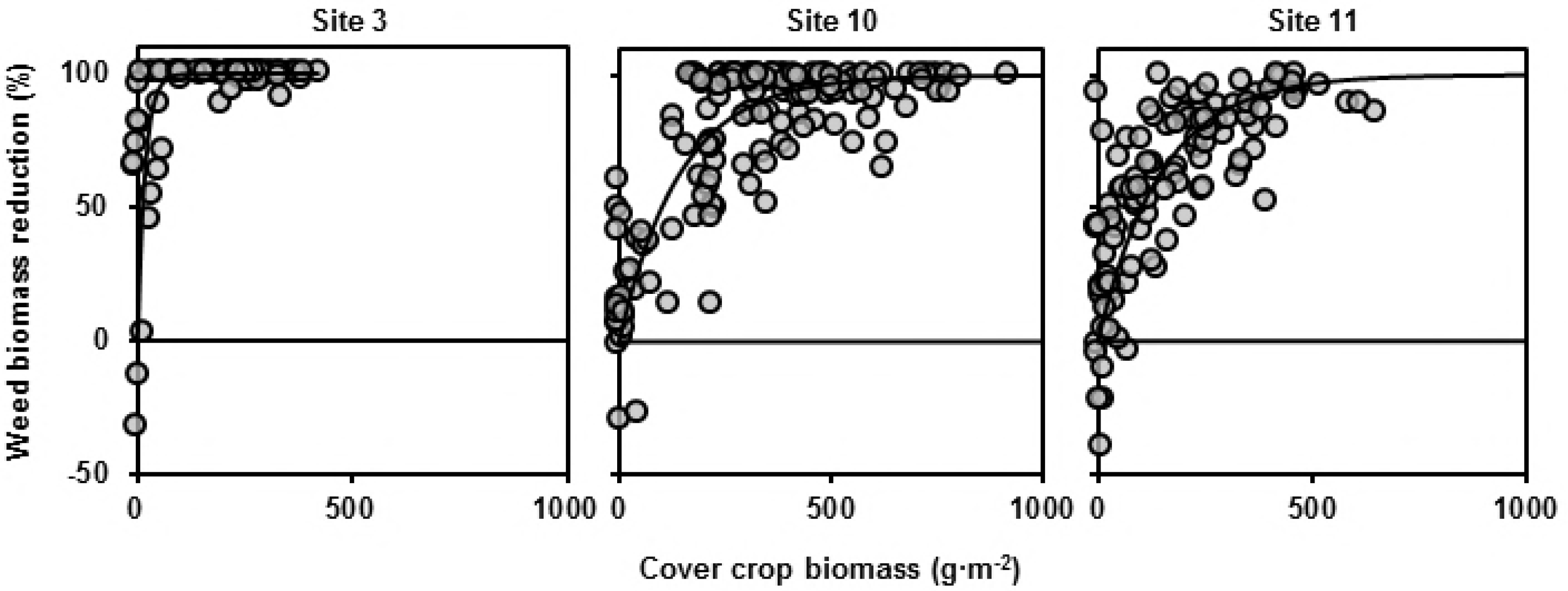
Effect of cover crop productivity on weed biomass reduction. Weed biomass reduction versus cover crop biomass at each of three sites. Exponential equation (Table 5) fit through each of the three data sets. Three data points with cover crop biomass beyond 1000 g m^−2^ not shown.

**Table 5.**
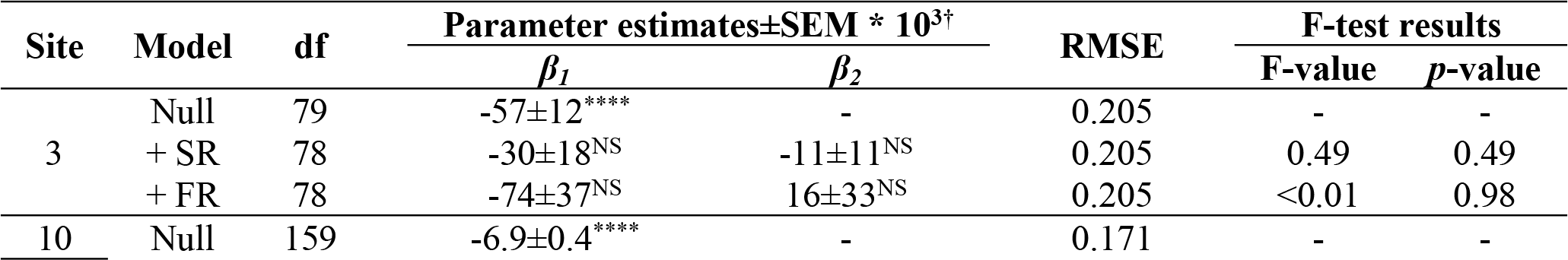
Parameter estimates for the exponential model fitted to weed biomass reduction versus cover crop biomass for each site with and without the inclusion of cover crop species richness (+SR) and functional richness (+FR) as a predictive variable along with F-test results. A significant value of ***β***_***2***_ indicates that cover crop diversity affects the relationship between weed biomass reduction and cover crop biomass.

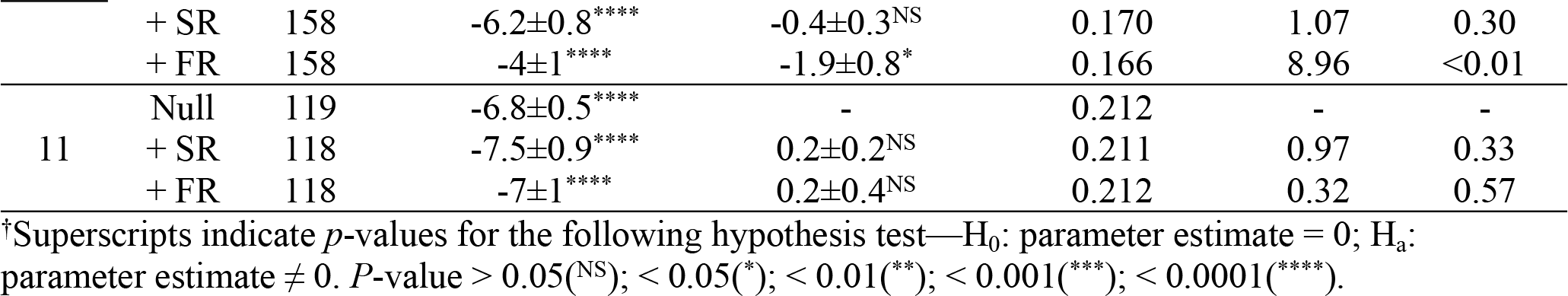

### Effect of diversity on stability

As mean cover crop biomass went up, so did the standard deviation (Fig 8). However, the slope of this relationship was not affected by cover crop mixture species richness or functional richness, suggesting that increasing cover crop mixture diversity does not stabilize biomass across individual sites (Table 6).

**Fig 8.**
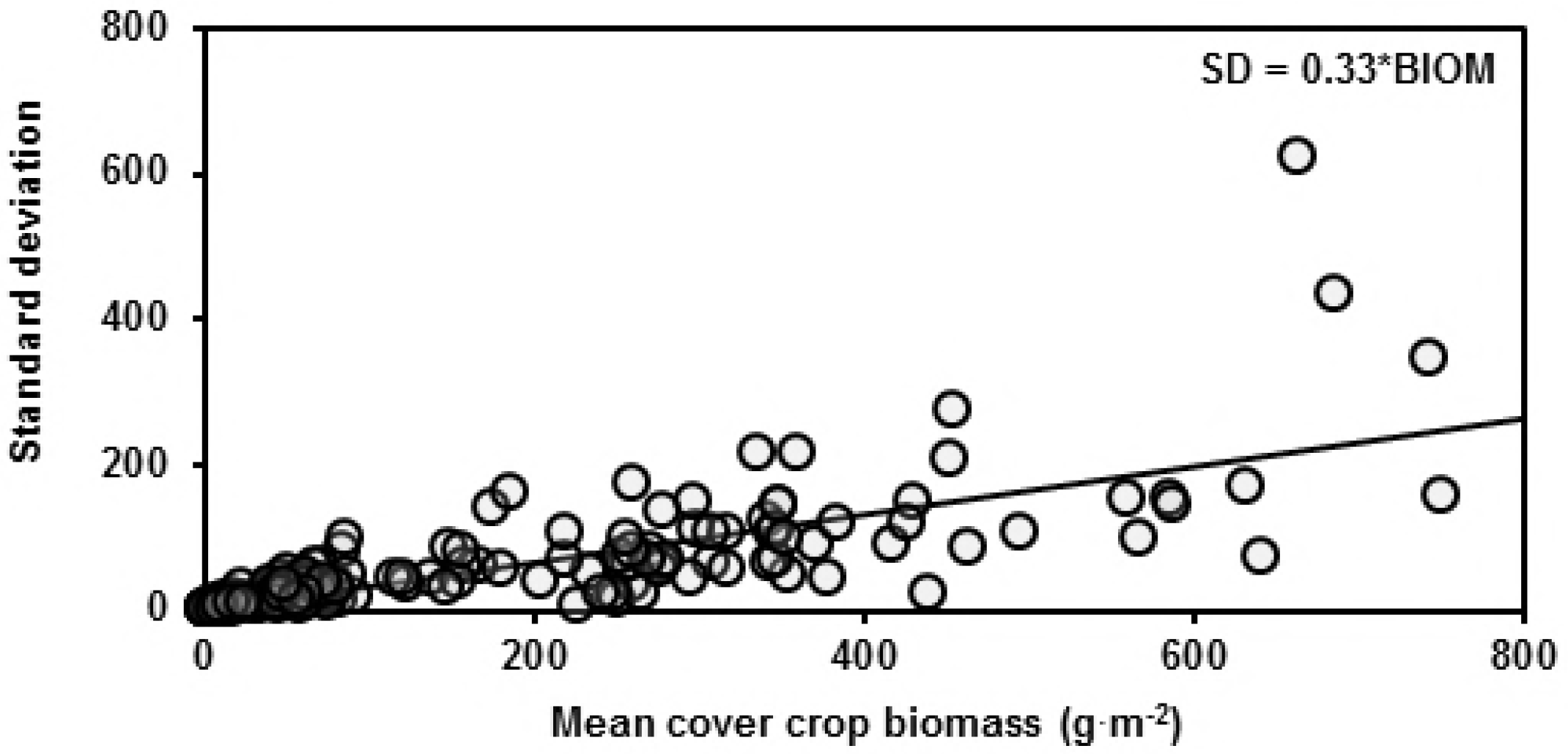
Stability of cover crop biomass. Standard deviation of cover crop aboveground biomass versus mean cover crop aboveground biomass for each treatment averaged across plots within each site. Line represents ordinary least squares regression with intercept term removed.

**Table 6.** Parameter estimates, degrees of freedom, and *p*-values for linear models relating standard deviation of cover crop biomass (SD) to mean cover crop aboveground biomass (BIOM) with and without cover crop species richness (SR) and functional richness (FR) interacting with cover crop aboveground biomass. A significant value of BIOM:SR or BIOM:FR indicates that species or functional richness, respectively, affects the relationship between SD and mean cover crop biomass.

| Equation<sup>†</sup> | df | Parameter<sup>‡</sup> | Estimate±SEM<sup>§</sup> | *p*-value |
| --- | --- | --- | --- | --- |
| SD ~ BIOM (Base model) | 172 | BIOM | 0.33±0.02**** | <0.0001 |
| SD ~ BIOM + BIOM:SR | 171 | BIOM | 0.35±0.02**** | <0.0001 |
|  |  | BIOM:SR | -0.006±0.005<sup>NS</sup> | 0.23 |
| SD ~ BIOM + BIOM:FR | 171 | BIOM | 0.38±0.03**** | <0.0001 |
|  |  | BIOM:FR | -0.03±0.01<sup>NS</sup> | 0.07 |

<sup>†</sup>Standard deviations and mean biomass determined for each treatment across plots within each site.
<sup>‡</sup>Intercepts fixed to zero.
<sup>§</sup>Superscripts indicate *p*-values for the following hypothesis test—H<sub>0</sub>: slope = 0; H<sub>a</sub>: slope ≠ 0. *P*-value > 0.05(<sup>NS</sup>); < 0.05(\*); < 0.01(\*\*); < 0.001(\*\*\*); < 0.0001(\*\*\*\*).

## Discussion

### Diversity-productivity hypothesis

Increasing plant mixture diversity, particularly functional richness, was associated with increased average aboveground biomass. However, at no site did the most productive mixture produce more biomass than the most productive monoculture. Both of these observations are consistent with the findings of the large majority of previous plant mixture studies in both agriculture and ecology [7, 14, 26–28].

The distinction between increasing average productivity and absolute productivity is not a trivial one. The diversity-productivity hypothesis asserts that increased diversity should lead to increased average productivity. It does so on the logic that diverse systems should have the potential to be more productive than even the most productive of monocultures by capturing a greater proportion of available resources [29]. That is, mixing plant species should be able to raise the ceiling on biomass productivity reached by plant monocultures. This, however, is a different conclusion than increasing diversity increases average productivity. According to the logic of niche complementarity, increasing diversity shouldn’t necessarily increase average productivity as is suggested by the diversity-productivity hypothesis. Rather, it should increase the absolute productivity. This disconnect between the theoretical underpinnings of the diversity-productivity hypothesis and the theoretical conclusions of the diversity-productivity hypothesis indicates (1) that we should be testing the theory of niche complementarity by testing whether increasing mixture diversity raises absolute productivity rather than average productivity and (2) that niche complementarity is not the necessary conclusion to be drawn from the observation that increasing diversity increases average productivity.

If we cannot ascribe our observation that increasing plant mixture diversity is associated with increased average productivity to increased niche complementarity or resource use efficiency, to what then can we ascribe this observation? The positive effect of increasing plant mixture diversity on average productivity is easily explained by low yielding species pulling down the average at low levels of diversity but not at high levels of diversity. Specifically, the pattern observed was simply the consequence of the average productivity of the monocultures and low functional richness category being brought down by the low yields of the legumes. In the high diversity treatments, the high yields of grasses and brassicas compensated for the low yields of legumes. This is why mixing across functional groups led to increased average productivity but not mixing within a single functional group. Mixing the grasses or the brassicas with each other did not increase average productivity because there were no low yielding species being compensated for in the mixture. Similarly, mixing the legumes together did not increase average productivity because there was no high yielding species in the mix to compensate for the low yields of the legumes.

Simply put, when there is bare space on the ground left by an unproductive species and we add another species, we get more vegetation. While this may seem like a simple description of niche complementarity, consider the fact that we could also get more vegetation by adding more of the same species. He et al. [30] found that the positive relationship between diversity and productivity decreased with increasing plant density.

### Diversity-invasibility hypothesis

A typical approach to evaluating the diversity-invasibility relationship is to evaluate an invasion resistance metric—e.g., weed biomass reduction—as a function of a diversity metric—e.g., cover crop species richness [27, 31–35]. Any positive trending relationship is then presented as evidence in favor of the diversity-invasibility hypothesis. The problem with this approach is that it mistakes correlation with causation, and confounds the effects of diversity with the effects of biomass productivity.

For example, analyzing our own data in this way, we found that weed suppression is positively correlated with cover crop species richness (Fig 9). However, since cover crop aboveground biomass was also correlated with species richness in this study (Fig 2), it is possible that the correlations between weed suppression and species richness were due to cover crop biomass rather than species richness. To determine whether or not species richness had an effect on weed suppression beyond its relationship with cover crop biomass, it was necessary to first control for the well-documented effect of cover crop productivity on weed suppression [36–37]. Controlling for the positive effect of cover crop biomass on weed suppression, we found that the effect of cover crop mixture diversity diminished markedly if it did not disappear entirely (Table 5).

**Fig 9.**
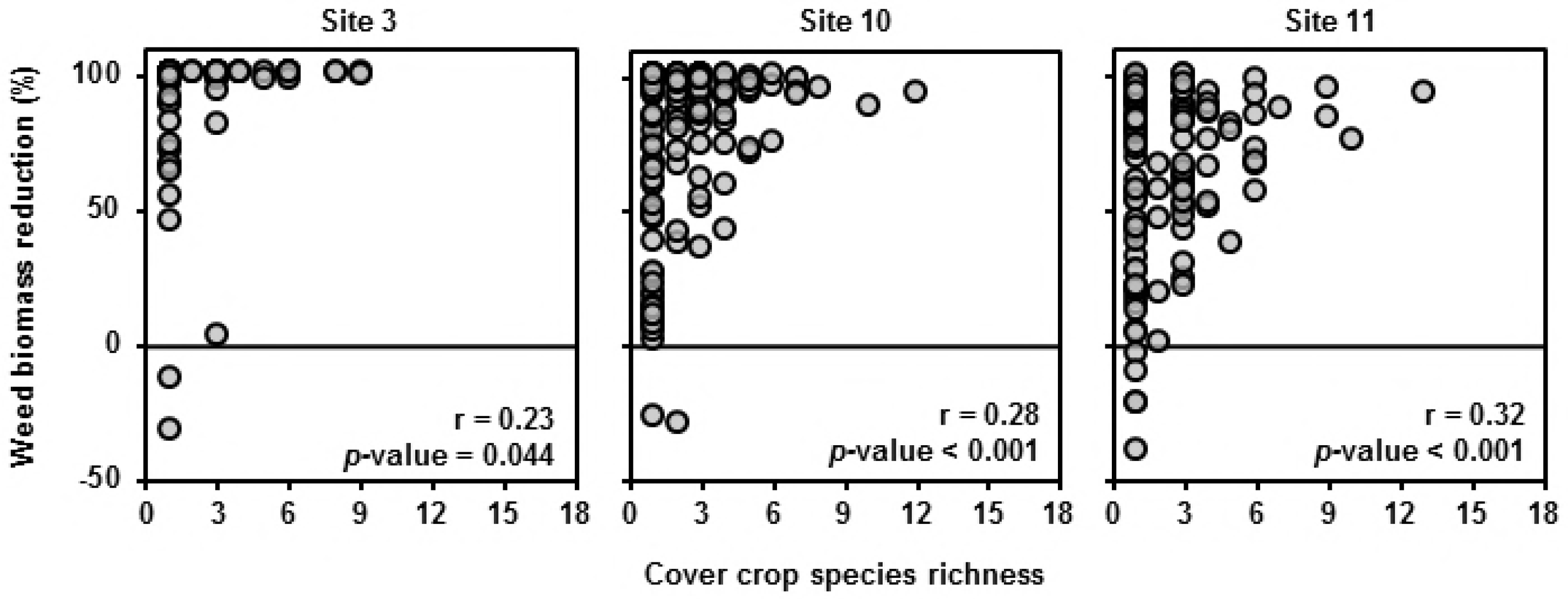
Effect of species diversity on weed biomass reduction. Weed biomass reduction versus realized cover crop species richness with Pearson correlation coefficients (r) for each site. *P*-values are for the following hypothesis test regarding the correlation coefficients—H_0_: r = 0; H_a_: r ≠ 0.

**Fig 2.**
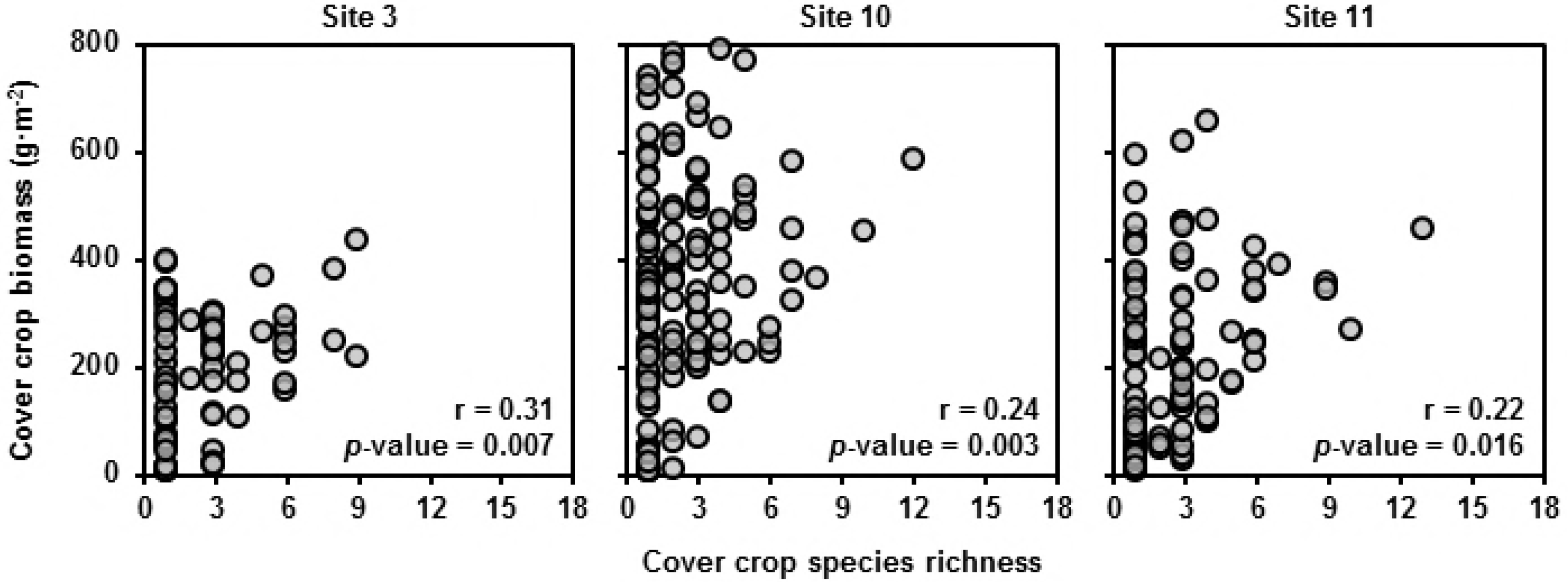
Cover crop biomass on species richness. Cover crop biomass versus realized cover crop species richness with Pearson correlation coefficients (r) for each site. Three data points with cover crop biomass beyond 1000 g m^−2^ not shown. *P*-values are for the following hypothesis test regarding the correlation coefficients—H_0_: r = 0; H_a_: r ≠ 0.

In most manipulated plant diversity studies, as plant diversity increases so does average biomass productivity [20, 29]. Increased plant productivity is well documented to be associated with increased invasion resistance in native systems and increased weed suppression in agricultural systems [38–41]. Yet most diversity-invasibility studies treat the correlation between plant mixture diversity and invasion resistance as evidence for the diversity-invasibility hypothesis, ignoring the mediating effects of biomass productivity on invader suppression, which may be driving much of the apparent effects of diversity on invader suppression. Subsequent meta-analyses that consolidate the findings of these studies also fail to address the confounding effects of biomass productivity on invader suppression [42, 43]. In the few studies where productivity is accounted for, the apparent effects of diversity on invasibility disappear [32, 44].

Reviews of mixed cropping literature give the impression that it’s the actual mixing of crops that is promoting weed suppression [45–47] without addressing the possibility that it could simply be increased biomass that results in greater weed suppression. Furthermore, if we use the increased weed suppression of intercrops as evidence of increased resource use efficiency, what of the cases where the sole crops are more suppressive than the intercrops [41, 48]? Do those results indicate that sole crops are more resource use efficient than intercrops? No, in order to explain this seeming inconsistency, we need to simply look at variations in biomass—sole crops that are more weed suppressive than intercrops tend also to be more productive in terms of biomass [41].

Our study highlights one of the major issues underlying most of the supposed evidence in favor of the diversity-invasibility hypothesis—the covariance of diversity with productivity. Goldberg and Werner [49] made an early call for scientists to account for the effects of biomass when studying plant invasion, but overwhelmingly their advice has been ignored. After accounting for the well-documented effect of plant productivity on weed suppression in this study, we observed little effect of cover crop diversity on invasibility (Table 5).

### Diversity-stability hypothesis

In diversity-stability studies, the most common approach to evaluating the effect of diversity on stability is to regress the coefficient of variation of stand biomass (Ĉ_v_) against a diversity metric—most often species richness (Fig 11) [22, 50–51]. We disagree with this approach because diversity co-varies with biomass productivity in our study and Ĉ_v_ is sensitive to biomass productivity (Fig 12). Consequently, the results of simply regressing Ĉ_v_ against diversity can be misleading because the effects of diversity on stability are confounded with the relationship between biomass productivity and stability.

**Fig 3.**
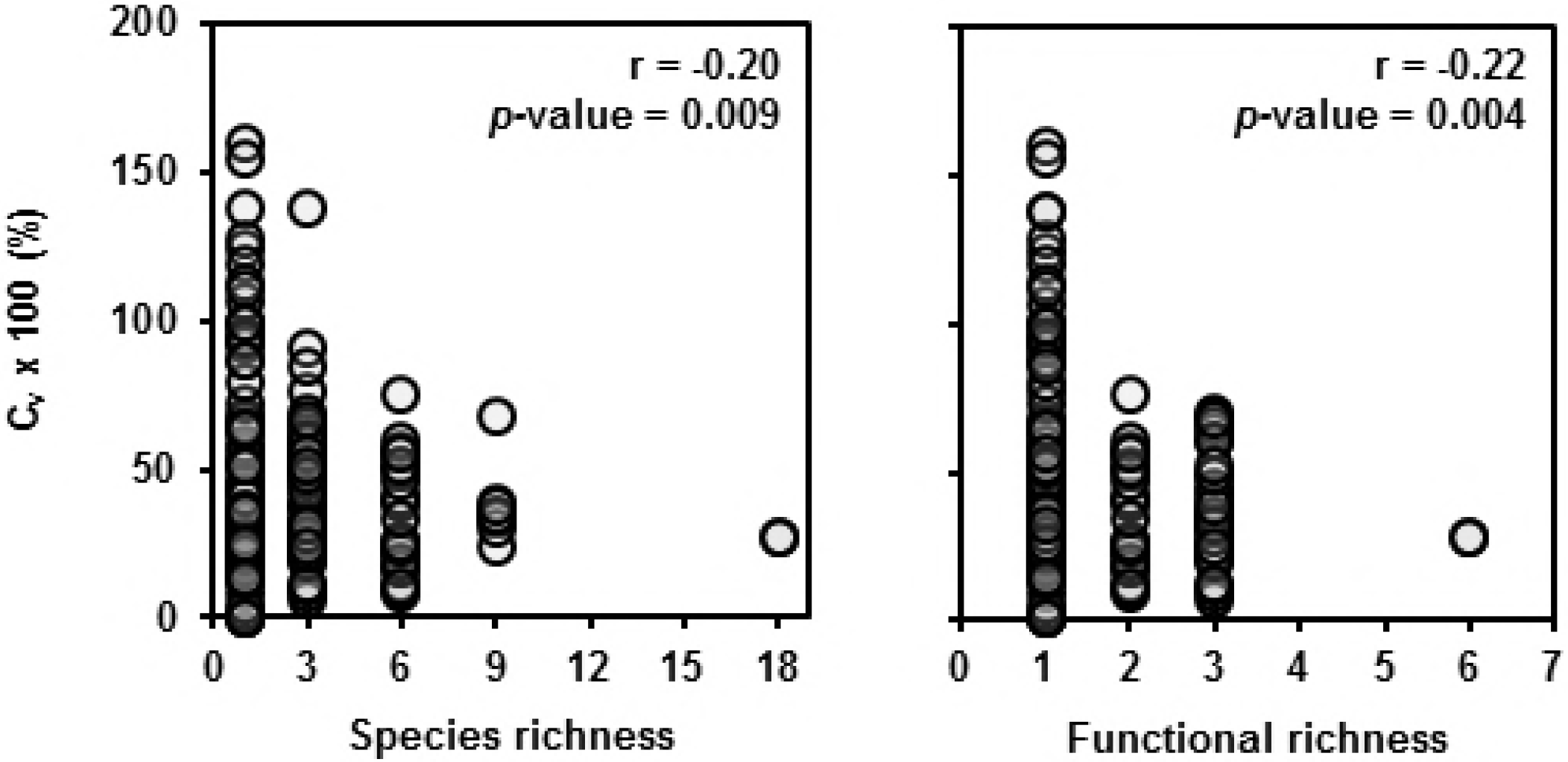
Effect of diversity measures on cover crop biomass coefficient of variation. Coefficient of variation of aboveground cover crop biomass across treatments and study sites plotted against realized species (left) and functional (right) richness. Pearson correlation coefficients (r) given with *p*-values for the following test—H_0_: r = 0; H_a_: r ≠ 0.

**Fig 4.**
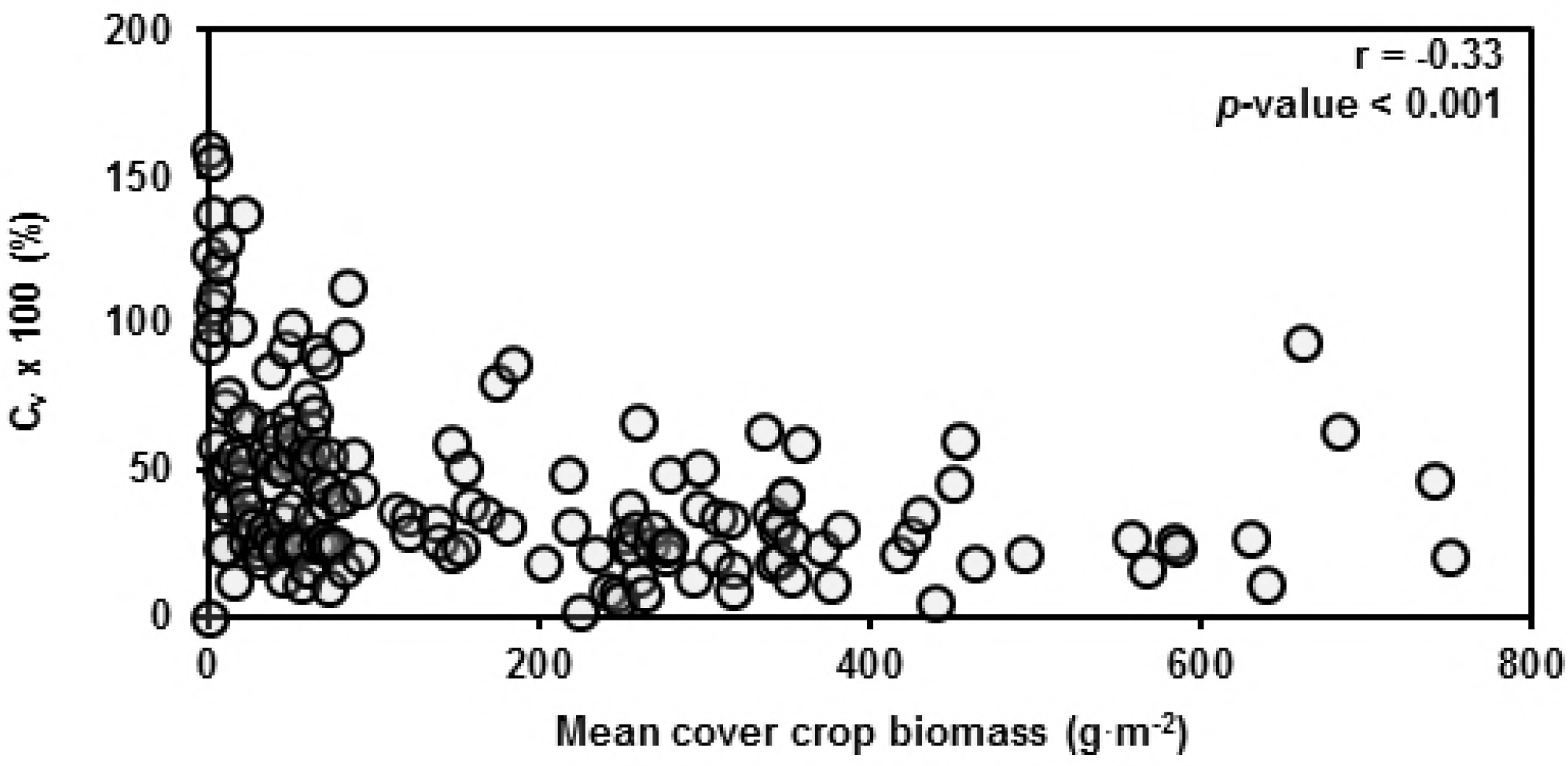
Cover crop coefficient of variation in relation to cover crop biomass. Coefficient of variation of aboveground cover crop biomass across treatments and study sites plotted against mean cover crop biomass. Pearson correlation coefficients (r) also given with *p*-values for the following test—H_0_: r = 0; H_a_: r ≠ 0.

Since C_v_ is calculated by dividing the standard deviation of a treatment by its mean biomass productivity, it would seem that the productivity effects are inherently accounted for in C_v_ calculations. The issue is not with the mean itself but rather with the interaction of the mean and the standard deviation. The Ĉ_v_ values were relatively constant beyond a certain level of mean biomass. At low levels of mean biomass, however, Ĉ_v_ values were unstable, which meant that less productive treatments on average had higher Ĉ_v_ values than more productive treatments (Fig 12).

To show how simply plotting Ĉ_v_ against diversity metrics might be misleading, we have done so with our data (Fig 3). Ĉ_v_ does decrease with increasing species and functional richness, but that’s not to say that increasing species and functional richness increases stability. If we look at the relationship between Ĉ_v_ and mean cover crop biomass, we find that at low biomass, the Ĉ_v_ tends to be greater and less consistent than at larger biomass (Fig 4).

These results occur because small amounts of experimental error at high levels of mean biomass have marginal effects on Ĉ_v_, whereas at low levels of mean biomass, small amounts of error amplify into dramatic effects on Ĉ_v_. Thus, the pattern that we observed in Fig 3 could simply have been because low diversity treatments tended to have less biomass in our study and treatments with less biomass tend to have higher Ĉ_v_. Increased species and functional richness were correlated with increased stability as measured by decreased Ĉ_v_ values. However, most of this effect was mediated by the covariance of diversity with productivity (Table 6).

Multiple intercropping studies have concluded that intercrops are more stable than sole crops on the basis of their C_v_ values being lower than those of the tested sole crops [52–53]. However, if we look at the productivity data of these studies, we find that the intercrops were also more productive than the sole crops. Furthermore, in cover crop mixture studies where the most diverse mixture is not the most productive treatment, neither are they the most stable [14, 28]. The Ĉ_v_ is clearly sensitive to mean biomass, and yet the effects of biomass on stability are rarely addressed in diversity-stability studies.

For the purposes of cover crop management, we found little evidence that increasing cover crop mixture diversity increased field-scale biomass stability. If we had greater species differentiation between the 18 species used and greater environmental heterogeneity, we might have expected a greater impact of diversity on stability, but for the practical purposes of cover crop management, where our environmental conditions are relatively predictable and our suite of potential cover crops thrive and fail under relatively similar conditions, that point may be moot.

## Conclusions

While increasing cover crop mixture diversity was often associated with increasing average cover crop biomass productivity, we contest the traditional interpretation of this result as evidence of increased niche complementarity or resource use efficiency of diverse mixtures. We argue that increased niche complementarity or resource use efficiency of mixtures should be indicated by increased absolute productivity rather than average productivity, which we did not observe. Our results are simply explained by the fact that the average biomass of monocultures was drawn down by low yielding species that were compensated for in mixture by high yielding species. While cover crop mixture diversity was often positively related to metrics of invasion resistance and stability, we found these correlations to be driven largely by variation in cover crop biomass. Once we controlled for the confounding factor of cover crop biomass, we found little evidence that cover crop mixture diversity positively affects invasion resistance or biomass stability.

Taken altogether, monocultures could be just as productive as mixtures and productive monocultures were just as effective at suppressing weeds and performing with the same stability as productive mixtures.

## Acknowledgments

Many thanks to cooperating farmers Ben Schole, Mike Herman, Paul Swanson, Keith Berns, and Nate Schroeder. Thanks also to Darren Binder and Paul Jasa for technical support. This material is based upon work supported by North Central SARE [GNC13-182] and by the National Science Foundation Graduate Research Fellowship [DGE-1041000].

